# A molecular brain atlas reveals cellular shifts during the repair phase of stroke

**DOI:** 10.1101/2024.08.21.608971

**Authors:** Rebecca Z Weber, Beatriz Achón Buil, Nora H Rentsch, Allison Bosworth, Mingzi Zhang, Kassandra Kisler, Christian Tackenberg, Ruslan Rust

**Author notes:** **Correspondence:** Ruslan Rust, PhD Assistant Professor, The Zilkha Neurogenetic Institute Department of Physiology and Neuroscience, Keck School of Medicine of the University of Southern California 1501 San Pablo Street, Room 341, Los Angeles, CA 90089 X: @rust_ruslan.

## Abstract

Ischemic stroke triggers a cascade of pathological events that affect multiple cell types and often lead to incomplete functional recovery. Despite advances in single-cell technologies, the molecular and cellular responses that contribute to long-term post-stroke impairment remain poorly understood. To gain better insight into the underlying mechanisms, we generated a single-cell transcriptomic atlas from distinct brain regions using a mouse model of permanent focal ischemia at one month post-injury. Our findings reveal cell- and region-specific changes within the stroke-injured and peri-infarct brain tissue. For instance, GABAergic and glutamatergic neurons exhibited upregulated genes in signaling pathways involved in axon guidance and synaptic plasticity, and downregulated pathways associated with aerobic metabolism. Using cell-cell communication analysis, we identified increased strength in predicted interactions within stroke tissue among both neural and non-neural cells via signaling pathways such as those involving collagen, protein tyrosine phosphatase receptor, neuronal growth regulator, laminin, and several cell adhesion molecules. Furthermore, we found a strong correlation between mouse transcriptome responses after stroke and those observed in human nonfatal brain stroke lesions. Common molecular features were linked to inflammatory responses, extracellular matrix organization, and angiogenesis. Our findings provide a detailed resource for advancing our molecular understanding of stroke pathology and for discovering therapeutic targets in the repair phase of stroke recovery.

## Introduction

Stroke remains a leading cause of disability and death, affecting one in four adults in their lifetime^1,2^. Over half of stroke patients are left with permanent disabilities including partial paralysis and cognitive deficits due to the brain’s limited ability to regenerate damaged neural circuits. Post-stroke damage develops from a complex interplay of pathological processes that involve all major cellular components of the brain including neurons, glia cells, resident and infiltrating immune cells, blood vessels, and peri-vascular mural cells.^3^ Each cell type undergoes significant changes in response to stroke and the contributions of individual cell types to recovery remain not fully understood. Some cellular responses were associated with enhances recovery, while others may be detrimental^4–7^. These phenotypic alterations are often regulated at the transcriptional level.

Several single cell/nucleus RNA sequencing studies characterized the cellular heterogeneity in the healthy human and mouse brain^8–14^. Recent cellular atlases were generated in response to several neurodegenerative diseases^15–17^ and acute neurological injuries including spinal cord injury and stroke, supporting the functional plasticity of individual brain cells^18–21^. Some molecular stroke atlases focused characterization of single cell types e.g., immune cells, pericytes or vascular cells^18,19,22^, some used acute time points^22–24^ and none looked at a permanent focal ischemia mouse model.

Therefore, we performed snRNAseq on mouse brains one-month post-stroke using a mouse model that simulates permanent stroke conditions to (a) generate a molecular atlas of cell types from stroke-injured and peri-infarct regions, (b) infer molecular communication networks among individual cells and (c) compare these transcriptomic changes to those observed in human chronic, non-fatal brain stroke lesions. These findings enhance our mechanistic understanding of stroke repair and may improve therapeutic targeting during the transition phase of subacute to chronic stroke.

## Results

### Cellular and molecular profiles of adult intact, peri-infarct and stroke-injured brain

To molecularly profile the stroke-injured brain, we induced large cortical strokes in the sensorimotor cortex of C57BL/6J mice (**Fig. 1A**). All mice showed a severe ≈70% reduction in cerebral blood flow in the stroke core, and ≈45% reduction in the ischemic border zone (ibz) compared to the intact hemisphere at 2 h after stroke induction (**Fig. 1B, C**).

**Figure 1:**
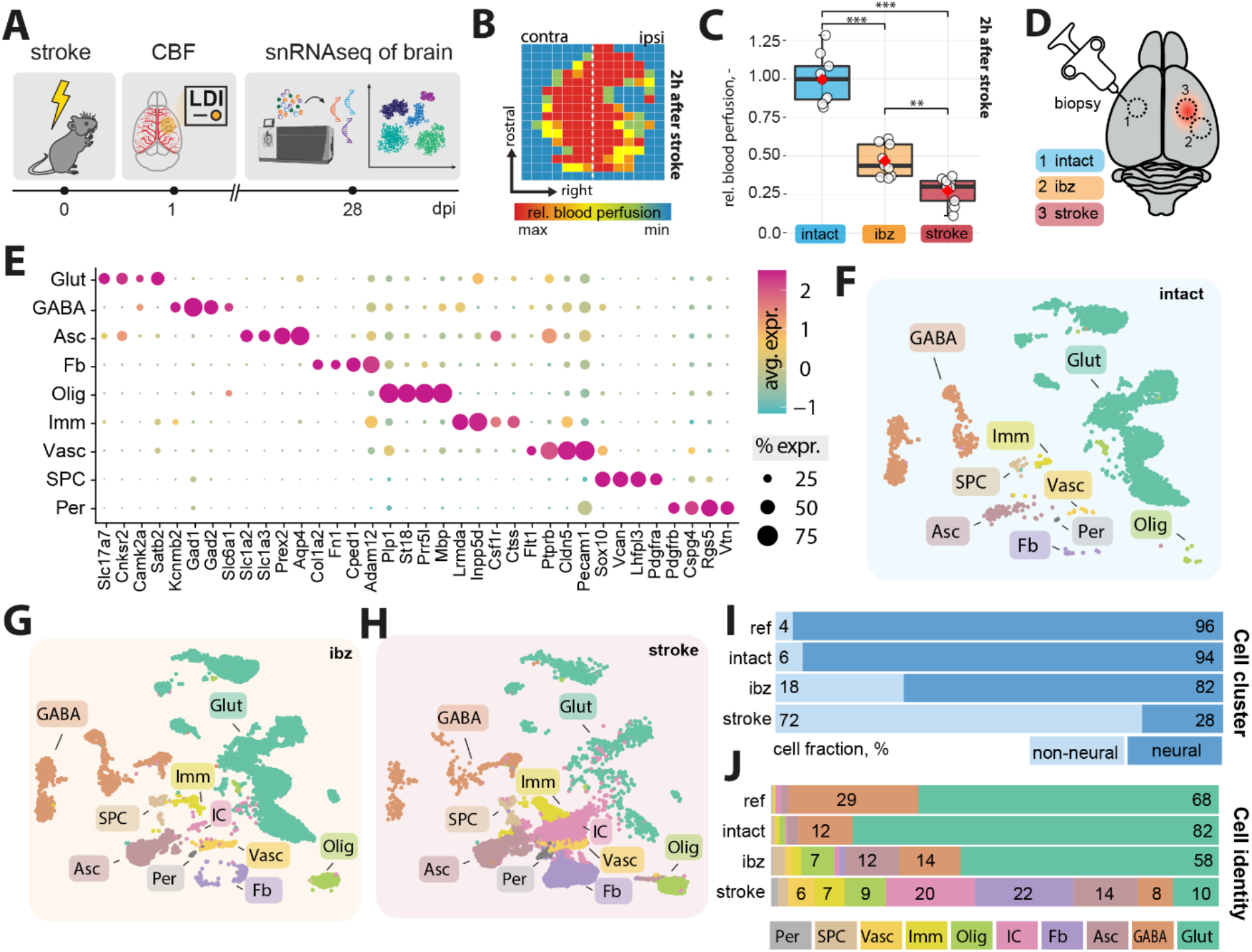
Cellular profiling of the stroke-injured mouse brain. (A) Scheme of experimental workflow (B) Laser Doppler imaging (LDI) illustrating relative perfusion in the mouse brain (C) Bar plot showing quantification of relative blood perfusion in the stroke core and the ischemic border zone (ibz) compared to the left hemisphere acutely after stroke (D) Illustration of brain regions for biopsy to perform snRNAseq (E) Dot plot representation of canonical cell type markers across different cell populations from the intact contralesional hemisphere, labeled by cell type: glutamatergic neurons (Glut), GABAergic neurons (GABA), astrocytes (Asc), fibroblasts (FB), oligodendrocytes (Olig), immune cells (Imm), vascular cells (Vasc), stem/progenitor cells (SPC), and mural cells (Per) (F-H) UMAP visualization of cell clusters from intact, ibz and stroke tissue. (I) Bar plot showing distribution of cell type by neural and non-neural cells and individual cell types (J) across the reference, intact, ibz and stroke samples. The data was generated with a cohort of n = 9 mice.

Four weeks after stroke, we performed a microbiopsy of a) intact, b) ibz, and c) stroke core tissue. All samples were processed for single nucleus RNA sequencing (snRNAseq) using the 10X Genomics Chromium platform, generating transcriptomes from approximately 35,000 nuclei (**Fig 1D, Suppl. Fig. 1**).

We performed clustering and annotation of nine major cell populations of the mouse brain guided by the known marker expression patterns from molecular atlases^11,17,25^ : Glutamatergic neurons (Glut: e.g., Slc17a7, Satb2), GABAergic neurons (GABA: e.g. Gad1, Gad2), astrocytes (Asc: Slc1a2, Slc1a3), fibroblasts (FB: Col1a, Fn1), oligodendrocytes (Olig: Mbp, Plp1), immune cells (Imm: Inpp5d, Csf1r), vascular cells (Vasc: Flt1, Cldn5), stem and progenitor cells (SPC: Sox10, Vcan), and mural cells (Per: Pdgfrb, Cspg4) (**Fig. 1E**). These cell types and marker expression of cell types matched previous single-cell/snRNAseq data from adult non-injured mouse cortices^12,26,27^.

We used these cell-type categories from the intact adult mouse brain to characterize changes specific to ibz and stroke-injured brain tissue (**Fig. 1F-H**). All cell types were determined with a high confidence in all datasets (**Suppl. Fig. 2**). We observe an increase in the proportion of non-neural cells in the ibz (+12%) and in the stroke-injured tissue (+66%, **Fig. 1I**). Notably, the ratio of both glutamatergic and GABAergic cells to the total cell population was reduced in the stroke-injured tissue (Glut: intact: 82%, ibz: 58%, stroke: 10%: GABA: intact: 12%, ibz: 14%, stroke: 8%), whereas the relative number of certain non-neural cell types increased especially e.g., fibroblasts (intact: 1%, ibz: 1%, stroke: 22%), astrocytes (intact: 4%, ibz: 12%, stroke: 14%), oligodendrocytes (intact: 2%, ibz: 7%, stroke: 9%), vascular cells (intact:1%, ibz: 3%, stroke: 6%) and immune cells (intact:1%, ibz: 2%, stroke: 7%, **Fig. 1J**).

### A distinct cell cluster termed injury-associated (IC) cells revealed by snRNAseq and immunohistochemistry

Next, we aimed to confirm the abundance of major cell populations in immunohistochemical stainings of intact and stroke brain tissue. We selected markers specific to mature neurons (NeuN^+^) and synapses (GAT^+^, vGlut1^+^, Synaptophysin^+^), astrocytes (GFAP^+^), macrophages (CD68^+^), microglia (Iba-1^+^), endothelial cells (CD31^+^) and pericytes (CD13^+^) and stained stroked and intact coronal brain sections 28 days after injury (**Fig. 2A, Suppl. Fig. 3**). We found that relative NeuN^+^ expression was significantly reduced in stroke-injured and ibz tissue, whereas GFAP^+^-expression increased in the ibz compared to the intact side (**Fig. 2B**). CD68^+^, highly expressed in macrophages^28^, and IBA1^+^, expressed in microglia/monocytes^23^, was found to be elevated in the stroke core and the ibz after injury, compared to marker expression in intact tissue (**Fig. 2B**). Interestingly, we also found increase in CD13^+^ signals in the stroke core that were not associated with CD31^+^ vasculature, potentially indicating recently described CD13^+^ infiltrating monocytes which have been described to aggravate acute stroke injure but promote chronic post-stroke recovery (**Fig. 2B**).^29^ Although the relative number of vascular cells, compared to other cell types, increased in the stroke snRNAseq dataset, the overall coverage of CD31^+^ vasculature in stroke tissue is lower in the injured hemisphere compared to the intact hemisphere, consistent with previous stroke studies^30–33^ (**Suppl. Fig. 3**). Furthermore, we found expression of platelet-derived growth factor receptor beta (PDGFR- β) to be upregulated in the stroke core and the ibz compared to intact tissue, as described before (**Fig. 2C, D**).^34^ After CNS injury, microglia create a pro-inflammatory environment by releasing TNF; and this has been shown to enhance the expression of integrins.^35,36^ We found that integrin subunit alpha V (ITGAV) was strongly expressed within the ibz, but barely in the intact cortex (**Fig. 2E,F**).

**Figure 2:**
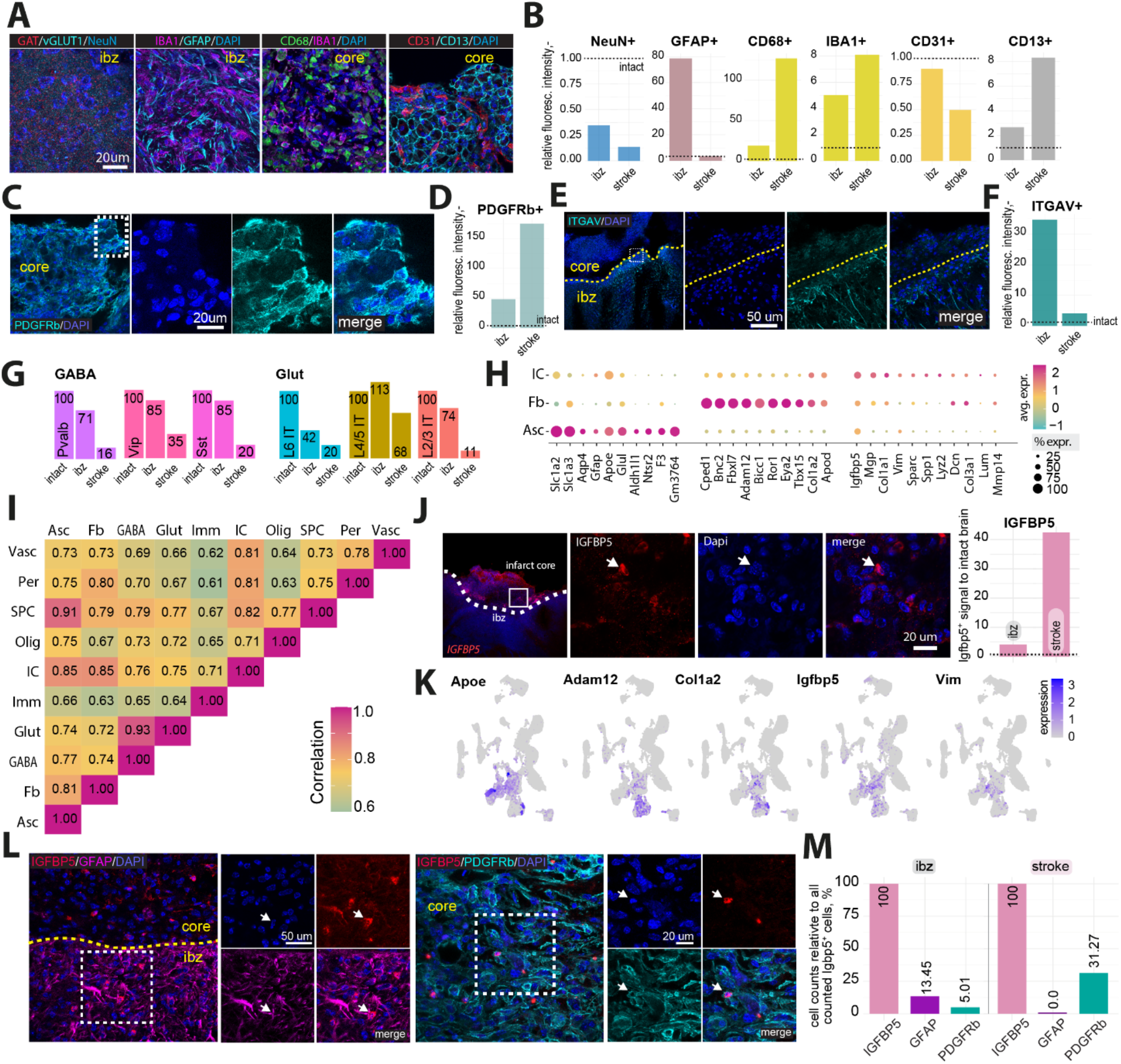
A distinct cell cluster termed injury-associated (IC) cells revealed by snRNAseq and immunohistochemistry. (A) Representative histological overview of brain sections stained with (from left to right) Gat/vGlut, IBA1/GFAP, CD68/IBA1, CD31/CD13, co-stained with DAPI. (B) Quantification of NeuN+, GFAP+, IBA1+, CD68+, CD31+ and CD13+ expression relative to intact tissue (dotted line). (C) Representative histological overview of brain sections stained with PDGFRb and co-stained with DAPI. (D) Quantification of PDGFRb+ expression relative to intact tissue (dotted line). (E) Representative histological overview of brain sections stained with ITGAV and co-stained with DAPI. (F) Quantification of ITGAV+ expression relative to intact tissue (dotted line). (G) Bar graph showing relative amount of major GABA and glutamatergic neuronal subtypes. (H) Dot plot showing expression of canonical cell type marker of injury-associated cells (IC), FB, and Asc. (I) Heatmap showing correlation of gene expression profiles between each cell type from stroke tissue. (J) Representative histological overview of brain sections stained with IGFBP5 (left) and quantification of IGFBP5 expression relative to intact tissue (right). (K) Feature plot showing the expression patterns of Apoe, Adam12, Cola1a2, and Vim in cells from stroke tissue. (L) Representative histological overview of brain sections stained with (from left to right) IGFBP5/GFAP and IGFBP/PDGFRB, co-stained with DAPI. (M) Cell counts (GFAP+ and PDGFRb+ positive cells, relative to all counted IGFBP5-positive cells) in the ibz and the stroke area. The data was generated with a cohort of n = 9 mice.

Analysis of neural subtypes using snRNAseq revealed a decrease in most glutamatergic and GABAergic subclusters in stroke-injured tissue (**Fig. 2G**). For instance, the ratio of GABAergic parvalbumin (PV)-expressing, vasoactive intestinal polypeptide (Vip)-expressing, and somatostatin (Sst)-expressing neurons, as well as glutamatergic layer 2/3 and layer 6 intratelencephalic (IT) neurons was reduced by >80% relative to the intact tissue. These findings align with previous functional studies showing that the loss of specific interneurons, such as PV and SST-expressing neurons, can worsen stroke outcomes and rescuing these populations may serve as a therapeutic target^37–40^.

Additionally, within the stroke core tissue, we identified a distinct cluster of cells, absent in the intact tissue, which we have termed ’injury-associated cells’ (IC) (**Fig. 1I, J**). IC cells are positive for astrocytic markers such as Apoe and Slc1a2 but also express genes related to ECM modeling such as Col1a2 and Col3a1. ICs transcriptionally segregated from other clusters by expression of genes involved in formation of scar tissue e.g., Dcn, Lum, Col3a1, and Col1a1, but also promotion of remodeling and tissue repair e.g., Mmp14, Vim, Igfbp5, and Sparc. Correlation analysis between all cell types revealed that the IC cell cluster shows most gene expression similarities (r = 0.85) to astrocytes and fibroblasts (**Fig. 2I, Suppl. Fig. 4**). Next, we selected one of the top IC markers and stained intact and stroked coronal brain sections (**Fig. 2J, K**). We found that insulin like growth factor binding protein 5 (Igfbp5) was strongly expressed within the infarction core, but barely in the intact hemisphere (**Fig. 2J, K**). To further characterize the IC cell cluster, we co-stained IGFBP5-positive cells with GFAP (reactive astrocyte marker) and PDGFR-β (fibrotic fibroblast-like cell marker)^41^ (**Fig. 2L**). We observed co-localization of PDGFR-β and IGFBP5 in both the infarction core (31% of all IGFBP5^+^ cells) and the ibz (5% of all IGFBP5^+^ cells), suggesting that these injury cells possess a fibrotic fibroblast-like identity (**Fig. 2M**). Additionally, 13.5% of IGFBP5^+^ cells in the border zone co-expressed GFAP, indicating that a subset of injury-associated cells may possess a more reactive astrocyte-like identity. Notably, ICs did also not show typical expression of canonical markers associated with fibrotic pericytes^4^.

Hence, these findings suggest that the IC cluster consists of cells with fibroblast-like and glial features that become reactive in response to CNS injury, consistent with previously observed cellular shifts in response to CNS injury^42–45^.

### Transcriptomic shift and pathway enrichment in brain cells following stroke

Next, we examined how long-term cerebral ischemia affects gene expression and pathway enrichment in individual brain cells (**Fig. 3A**). We found that most cell types exhibited differentially expressed genes (DEG) after stroke. As expected, most DEG were observed between nuclei from stroke-injured brains compared to intact tissue. Most DEGs were observed in FB, Glut and GABA nuclei (**Fig. 3A**). Interestingly, most DEGs overlapped between stroke/intact and ibz/intact tissue for neural (Glut and GABA) nuclei (**Fig. 3B**), whereas non-neural cells exhibited a more distinct, regional-specific DEG signature (**Fig. 3C**).

**Figure 3:**
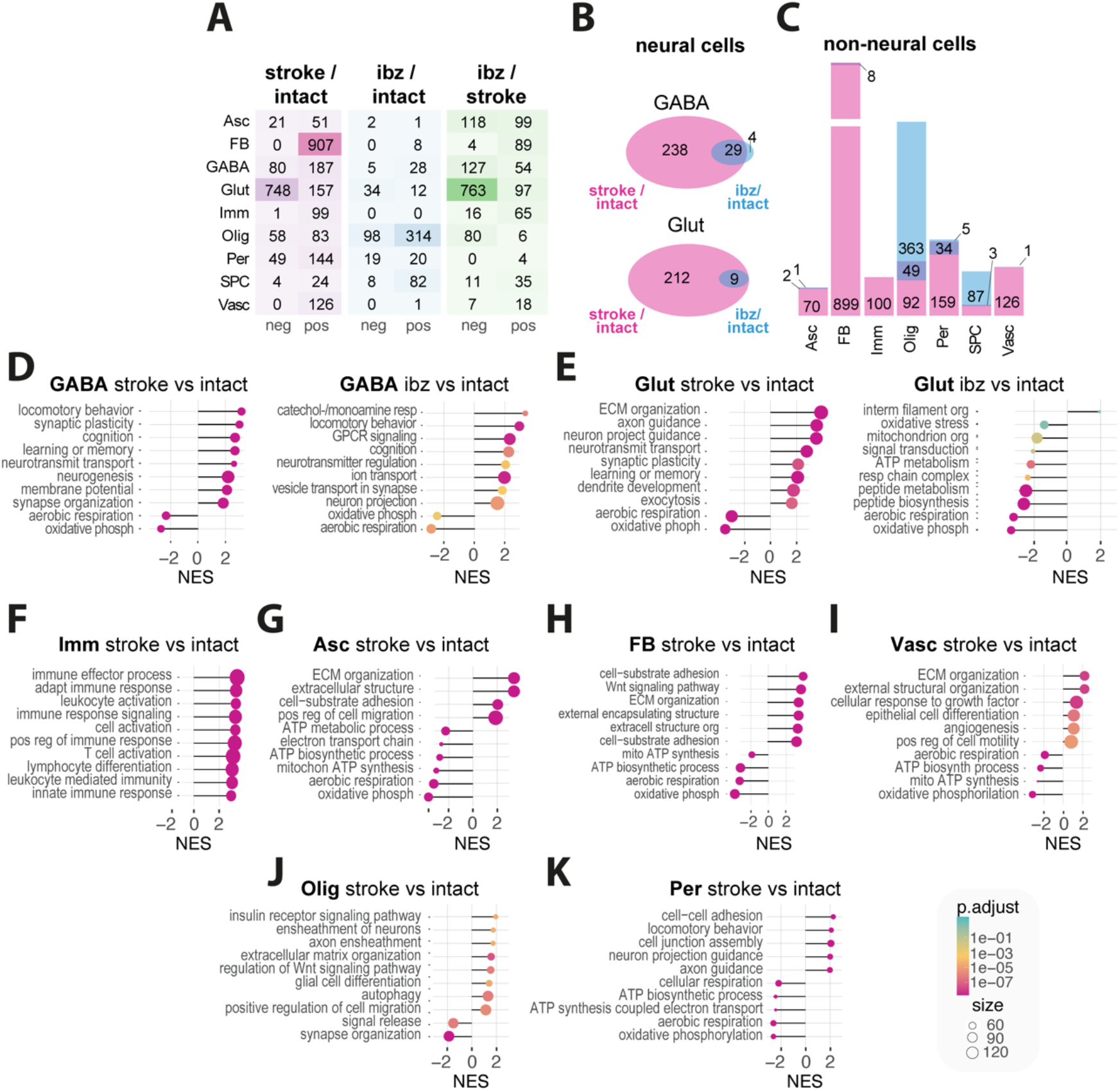
Transcriptomic responses of individual cell types to stroke in distinct mouse brain regions. (A) Heatmap showing number of significantly up- and downregulated genes per cell types in stroke vs intact (left), ibz vs intact (middle) and ibz vs stroke (right) tissue. (B) Venn diagram showing the overlap and unique differentially expressed genes from neural cells between stroke and ibz tissue (C) Bar plot showing the common and differential expressed genes in non-neural cells (right). (D-I) Gene set enrichment analysis (GSEA) of biological pathways that are enriched in stroke vs intact and ibz vs intact tissue for (D) GABA, (E) Glut, and GSEA from stroke vs intact tissue in (F) Imm, (G) Asc, (H) FB, and (I) Vasc (J) Olig, and (K) Per. Each panel displays the normalized enrichment score (NES) for pathways that are overrepresented (positive NES) or underrepresented (negative NES) in the post-stroke environment compared to intact tissue.

Gene set-enrichment analysis (GSEA) revealed that upregulated pathways in GABA and Glut neural cells predominantly involved pro-regenerative responses including synaptic plasticity, neurotransmitter transport, synapse organization, and axon guidance (all p < 0.001), while downregulated pathways in all neural cells included aerobic respiration and oxidative phosphorylation (**Fig. 3D, E**; all p < 0.001). This shift in cellular metabolism from energy-efficient aerobic respiration to alternative metabolic processes may potentially reflect an adaptive response to the altered microenvironment post-stroke^46^.

Immune cells mainly showed an upregulation in inflammation-associated pathways such as leukocyte activation and positive regulation of immune response (all p < 0.001). Notably, immune cells were the only cell types that did not exhibit altered aerobic metabolism (**Fig. 3F**). Astrocytes and fibroblasts revealed enrichment in pathways linked to remodeling such as extracellular matrix (ECM) organization and cell adhesion and migration processes (all p < 0.001) (**Fig. 3G, H**). Additionally, ECs showed enrichment in angiogenesis and remodeling pathways, alongside a downregulated in aerobic metabolism (**Fig. 3I**; all p < 0.001). Oligodendrocytes show significant enrichment in pathways related to axon ensheathment, extracellular matrix organization, and Wnt signaling regulation, suggesting an adaptive role in myelin remodeling and structural support post-stroke (all p < 0.001) (**Fig. 3J**). Pericytes revealed a significant enrichment in pathways related to cell adhesion and axon guidance and reduction in ATP biosynthesis and oxidative phosphorylation, indicating a metabolic adaptation to reduced (all p < 0.001) (Fig. 3K).

Together, these data suggest major transcriptional changes of all major brain cells at 28 days after stroke involving pro-regenerative and remodeling pathways, while also indicating a persistently inflammatory and hypoxic environment.

### Mapping of intercellular molecular communication after stroke

To quantitatively infer and identify relevant communication networks after stroke, we used CellChat^47,48^ to analyze signaling patterns involved in ligand-receptor interactions between individual cell types (**Fig. 4A**). Our analysis suggests that the total number and strength of predicted interactions are increased in stroke tissue (**Fig. 4B, C**; number of interactions: stroke / intact: +104%, stroke / ibz: +71%: interaction strength: stroke / intact: +95%; ibz / intact+104%).

**Figure 4:**
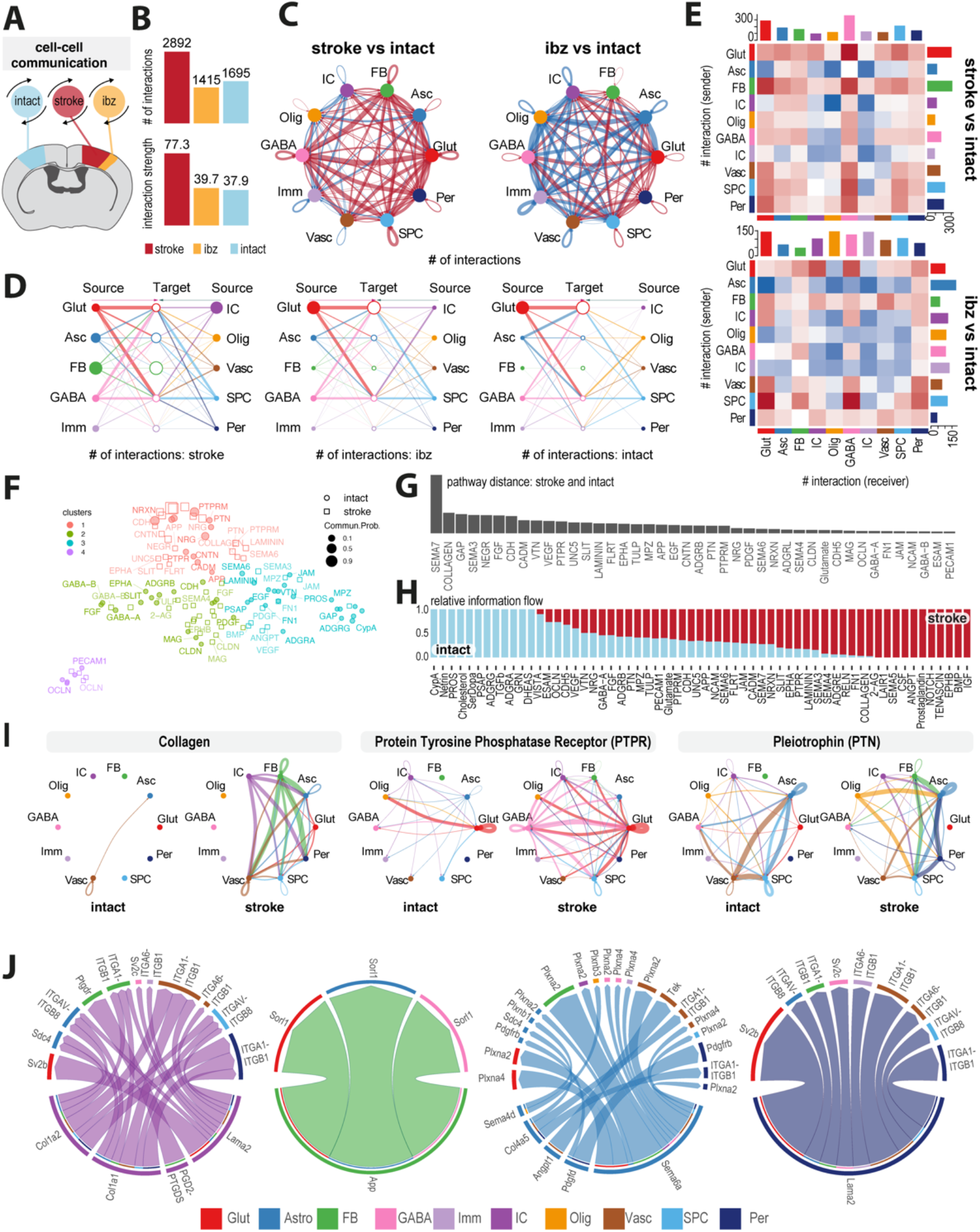
Mapping of intercellular molecular signaling post-stroke. (A) Schematic of cell-cell interaction analysis (B) Bar plot showing number and strength of interactions in cells from intact, ibz and stroke tissue. (C) Network diagram contrasting total number of cell-cell interactions between individual cell types in stroke vs intact (left) and ibz vs intact (right). Red lines indicate increased interaction, blue lines indicate reduced interaction, relative to intact tissue. (D) Hierarchy plot of interaction between all individual cell types to target cells in stroke (left), ibz (middle) and intact (right) datasets. (E) Heatmap showing differential interactions between cell types from stroke vs intact (upper) and ibz vs intact tissue (lower). Red squares indicating increased signaling and blue squared indicating decreased signaling, relative to cells from intact tissue. (F) Scatter plot projecting signaling groups onto a 2D space according to their functional similarity between cells from stroke and intact tissue. (G) Bar plot showing signaling pathway distance between stroke and intact tissue (H) Stacked bar plot illustrating the proportional relative information flow in signaling pathways between intact and stroke tissue. (I) Cell-cell communication networks for selected pathways: COLLAGEN, Protein Tyrosine Phosphatase Receptor (PTPR), and Pleiotrophin (PTN) across cell types in intact (left) and stroke (right) tissue. (J) Chord diagrams showing the most upregulated signaling ligand-receptor pairings in injury-associated cells (IC), fibroblasts (FB), astrocytes (Asc), and pericytes (Per).

In stroke-injured tissue, we observed an upregulation of interactions among individual cell types (**Fig. 4C**), with the majority of communication occurring between information sending Glut, GABA, Asc, IC and FB and information receiving Glut, GABA and Asc (**Fig. 4D**). By contrast, cells derived from the ibz and intact tissue exhibited a lower number of predicted interactions especially among non-neural cell populations such as Imm, Vasc, Per, IC and Olig compared to corresponding cell types from stroke-injured tissue (**Fig. 4D, E**). In stroke-affected tissue, Glut and Asc, particularly as senders, display significantly increased interactions, with IC also showing heightened communication compared to intact tissue. Conversely, in the ibz, these interactions are less pronounced, with non-neural cells like Imm and Vasc cells engaging in fewer communications overall. The data suggests a substantial upregulation of neural cell interactions post-stroke, with a notable contribution from IC cells in stroke conditions (**Fig. 4E**).

To better understand the involved signaling pathways in stroke compared to intact tissue, we grouped and clustered signaling pathways in four groups separated by functional similarity (**Fig. 4F**). Most divergent pathways were linked with important biological functions such as neuronal guidance and plasticity (SEMA3, SEMA7, UNC5, SLIT, EPHA), vascular repair and ECM remodeling (COLLAGEN, LAMININ, CDH, CADM, ANGPT, FGF, MMP9).

Most of these pathways demonstrated a considerably higher information flow in stroke-injured tissue (**Fig. 4G, H, Suppl. Fig. 5, 6**). For instance, stroke tissue showed enhanced communication via the COLLAGEN, PTPR, PTN, NEGR, LAMININ, CNTN and CADM pathways, involving multiple cell types that either did not participate or exhibited only minimal signaling interactions in intact tissue (**Fig. 4I, Suppl. Fig. 7**). Interactions of individual cell types in these pathways reveal that ICs preferentially signal to non-neural cells through networks related to COLLAGEN and LAMININ networks involving e.g., Col1a2, Col1a1, Lama2. These signals are predicted to be received by astrocytes (e.g., Itgav-Itgb8), fibroblasts (itga1-itgb1, Ptgdr) and vascular cells (Itga1-itgb1, Itga6-itgb1). This signaling pattern appears to be distinct from the signaling of other non-neural cells such as fibroblasts (e.g. App), astrocytes (e.g., Sema6a, Sema4d, Angpt1) and pericytes (e.g., Lama2) (**Fig. 4J**).

Overall, these findings show the surprisingly complex and dynamic communication among individual cell types in stroke-injured brains.

### Comparative analysis of transcriptome responses in mouse and human stroke

To decode and highlight transcriptomic changes that may be relevant to human stroke, we compared our mouse pseudo-bulk and ortholog-transformed RNAseq data with publicly available human stroke-lesion RNAseq datasets from cortical lesions and contralesional brain tissue of patients who experienced a nonfatal ischemic stroke up to five years before death (GSE56267)^49^ (**Fig. 5A**).

**Figure 5:**
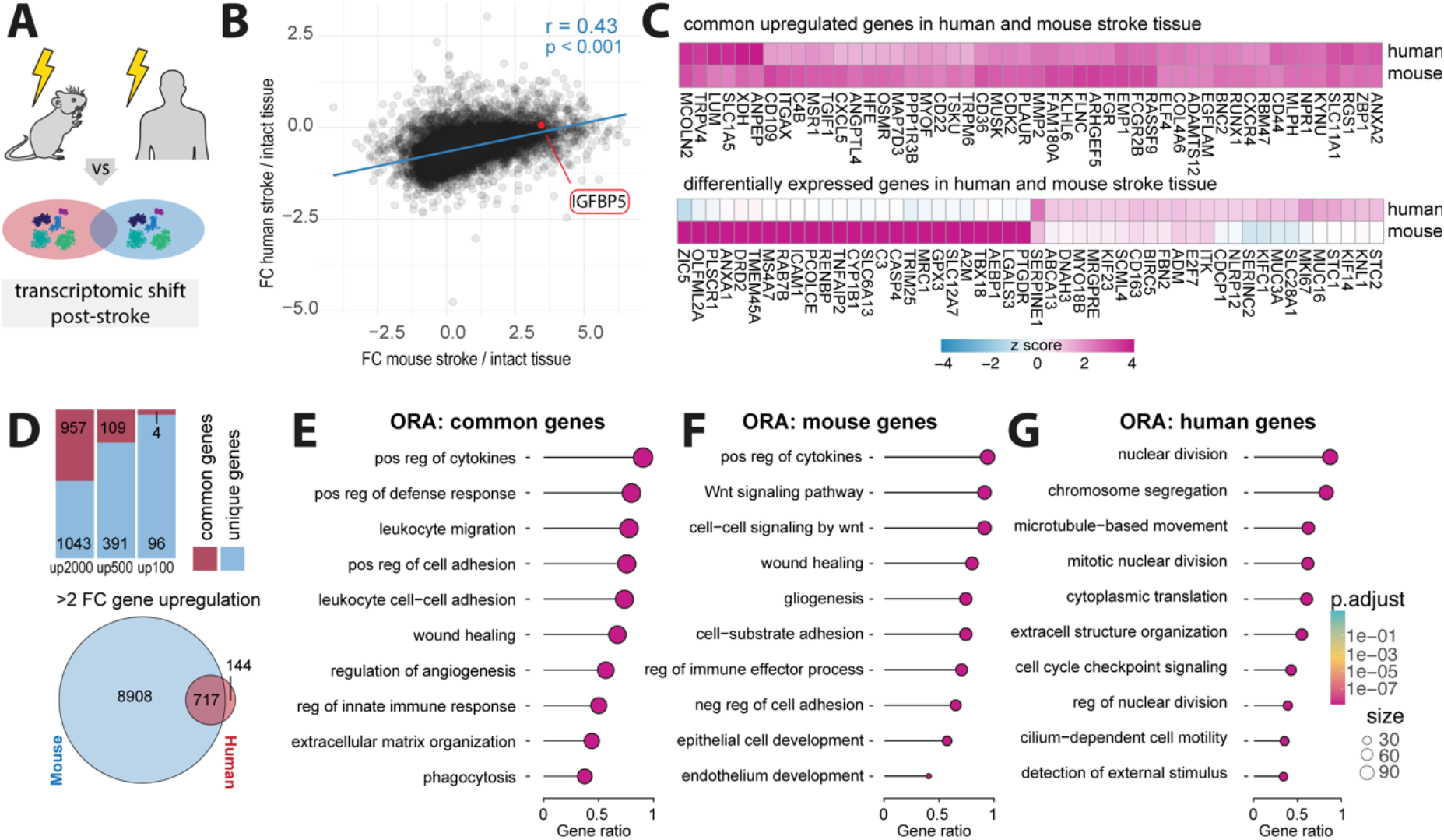
Comparative gene expression profiles in mouse and human post-stroke. (A) Schematic of human and mouse stroke (B) Scatter plot displaying the Person correlation for gene expression changes between mouse and human post-stroke. (C) Heatmap of common upregulated genes in human and mouse stroke (upper) and differentially expressed genes in human and mouse stroke (lower) (D) Stacked barplot of common and unique upregulated genes in the top 2000, 500 and 100 gene sets from both mouse and human datasets (upper) and Venn diagram of shared and unique genes with a fold change >2. (E-G) Overrepresentation analysis of top 2000 genes present in (E) both mouse and human datasets, (F) genes exclusively identified in mouse stroke dataset (G) and genes only present in human stroke dataset. All p-values ***,< 0.001.

We performed a Pearson correlation analysis of the genes shared by mouse and human datasets post-stroke, which revealed similarities in gene expression changes (r = 0.43, p < 0.001) (**Fig. 5B**). Interestingly, we found that IGFBP5, previously identified as upregulated in IC cells of stroke mice, was also upregulated in the human stroke dataset (**Fig. 5B**). We then calculated z-scores of log2-fold changes and revealed shared and differentially expressed genes between the mouse and human datasets (**Fig. 5C**). The overlapping upregulated gene expression featured genes associated with inflammation (e.g., CXCL5, CD44, CD36), neural plasticity (e.g., GAP43, RUNX1), ECM remodeling (e.g., ADAM12, MMP2, COL4A6) and vascular remodeling (e.g., ANGPTL4, CLDN5).

We observed that 48% of the top 2000 genes, 22% of the top 500 genes, and 4% of the top 100 genes were commonly upregulated in both mouse and human post-stroke. Moreover, of the 861 human genes exhibiting more than a 2-fold upregulation following stroke, 717 (83%) were also upregulated in the mouse stroke dataset (**Fig. 5D**).

We then conducted an over-representation analysis (ORA) of biological processes among the top 2000 upregulated genes that were (a) common: upregulated in mouse and human, (b) mouse-specific, and (c) human-specific. The common ORA predominately featured inflammation-related pathways including regulation of cytokines and immune cells activation, along with angiogenesis, ECM regulation and wound healing responses (**Fig. 5E**; all p < 0.001).

Mouse-specific ORAs similarly highlighted pathways of inflammation, wound healing, and gliogenesis (**Fig. 5F**; all p < 0.001). In contrast, human-specific ORAs were primarily associated with pathways involved in cell division and structural organization (**Fig. 5G**; all p < 0.001), potentially highlighting species-specific differences in the cellular repair mechanisms post-stroke.

In summary, our analysis underscores the substantial cross-species similarities in stroke-induced transcriptomic changes, especially in key pathways related to inflammation and tissue remodeling, highlighting conserved biological mechanisms that could inform the development of therapeutic strategies for human stroke recovery.

## Discussion

In this study, we have provided a resource for exploring single-cell transcriptomic data from distinct brain areas one month after stroke injury. Understanding the transcriptomic shifts of individual brain cells and the intracellular signaling mechanisms involved may further help to identify novel therapeutic targets. Our comparative analysis to human chronic stroke dataset has shown a considerable overlap in molecular gene and pathway enrichment suggesting similarities in the pathophysiology of chronic stroke between mice and humans.

Our findings indicate a considerable reduction in the glutamatergic and GABAergic neurons, which has been previously shown to affect stroke recovery^38,50^. Interestingly, gene expression profiles of the surviving cells, irrespective of their type, indicated a downregulation of genes involved in aerobic metabolism, implying a state of persistent hypoxia within the stroke regions. This observation aligns with previous studies that have documented prolonged alterations in cerebral blood flow post-stroke in both murine models and human patients^51,52^. While all neurons mainly rely on aerobic metabolism^53^, neurons with especially high energy demands, such as glutamatergic pyramidal neurons and fast-spiking GABAergic neurons (e.g., PV-expressing interneurons), may be even more susceptible to hypoxia^39^.

In glutamatergic neurons, we observed upregulation of axon guidance and synaptic organization pathways, partially mediated through semaphorin (Sema) signaling. Sema3E, which is highly expressed in the thalamus, and its receptor Plexin-D1 play a key role in neuronal circuit remodeling after stroke. By guiding synaptic connectivity, particularly in direct pathway medium spiny neurons (MSNs), Sema3E/Plexin-D1 signaling can help refine or restore motor and cognitive functions^54,55^. Various studies have reported an increase of guidance factors, including Sema3e after stroke^30,56,57^. There is evidence suggesting that blockage of Sema3e/PlexinD1 pathway might offer therapeutic benefits for restoring function after stroke^41^. However, other findings indicate that genetic deletion of PlexinD1 signaling can lead to impairments in the BBB^56^, thus, further studies are needed to clarify its dual roles in synaptic plasticity and vascular integrity.

In our study, we observed a pronounced upregulation of genes and pathways associated with ECM remodeling in non-neural cells including astrocytes, fibroblasts, and vascular cells. This upregulation is consistent with the well-known formation of a fibrotic scar in the stroke core, closely bordered by a glial scar after stroke. Importantly, we observed the ECM remodeling molecular signature in multiple non-neural cell types within the stroke, supporting the previously described cellular composition of the scar including ECM-producing PDGFRβ^+^ stromal fibroblasts, pericytes, reactive astrocytes, microglia, and monocyte-derived macrophages^41^. While the glial scar has been known for its dual role, potentially protecting the brain from further damage^58–60^, the fibrotic scar is primarily regarded as detrimental, particularly due to its inhibitory effect on regeneration^41,61^. Through cell-cell communication networks, we predicted that most upregulated signaling pathways include major components of ECM remodeling including collagen, integrin, and laminin pathways in multiple non-neural cells. Several integrin subunits α1β1, αvβ3, and α6β1, primarily expressed on endothelial cells, have been shown to regulate vascular remodeling and attenuate BBB permeability following stroke^62–64^. Several integrin subtypes have also been described to be involved in astrocyte scar formation and the phenotypic switch of astrocytes^65–67^. Notably, certain integrins, such as αIIbβ3, α4β7/α4β1, and αLβ2, are druggable targets that have been explored in cardiovascular diseases and inflammatory bowel disease^68^. Future research therefore may explore integrin-targeted therapies in stroke.

When comparing our dataset to scRNAseq datasets that were performed at earlier timepoints after stroke (3dpi)^69^, we found that several genes, including Lgals3 (Gal-3) and Serpina3n, display sustained upregulationin microglia beyond the acute post-stroke phase, suggesting that certain inflammatory and reparative pathways remain active through early and late stages of recovery. Gal-3 has been shown to increase in proinflammatory microglia in the early phase after ischemic stroke, where it promotes activation and proliferation of microglia.^70,71^ Notably, genetic deletion of Gal-3 attenuates cytokine release and neurodegeneration, highlighting its potential role in modulating neuroinflammation.^72^ Additionally, Gal-3 supports angiogenesis via VEGF-dependent mechanisms, which may extend into the chronic phase to facilitate tissue remodeling.^73,74^ Similarly, Serpina3n, a serine protease inhibitor enriched in astrocytes and microglia, can exert anti-inflammatory effects and help maintain BBB integrity following cerebral ischemia.^75^ Intranasal administration of recombinant Serpina3n protects against BBB breakdown and promotes long-term functional recovery when administered as early as 30 minutes post-stroke. Given our observation that Serpina3n remains elevated at 28 days post-injury, it would be important to investigate whether later therapeutic interventions could also exploit this pathway. Together, these data support the notion that targeting sustained neuroinflammatory and reparative processes, such as those involving Gal-3 and Serpina3n, might provide extended therapeutic windows and potentially improve long-term outcomes following ischemic stroke. Similarly, we also observed shared intercellular communication pathways including ANGPTL and SEMA3, which were previously shown to regulate acute and subacute vascular stability, cell migration, and neurovascular remodeling in response to CNS injury.^56,76^

We identified a cell cluster of cells, termed ‘injury associated cells’ (IC) in the stroke core that was barely present in peri-infarct nor intact tissue. There is evidence suggesting that ICs may be fibroblast-like cells that migrate to injury sites after stroke. However, their molecular characterization is challenging since there are only few fibroblast-specific markers and many of those markers can be substantially altered after injury^77^. For instance, we observed presence of Col1α1 and Vim, markers that have been used to identify fibroblasts, however, those markers can also be found on other cell types such as activated astrocytes^78,79^ or injury-associated pericyte-like cells (type A pericytes)^80^. The function of Col1α1+ associated cells in the injured brain has been described as both neuroprotective and detrimental.^63,64^ Studies have shown that Col1α1+ fibroblast-like cells are a crucial source of retinoic acid, promoting neural progenitor differentiation and improving recovery in rodent stroke models^80^.

Furthermore, research on chronic mouse ischemic stroke models reveals that Col1^+^ fibroblast-like cells in peri-infarct areas strongly express periostin, which has been linked to improved recovery in neonatal hypoxic-ischemic mice by promoting neural stem cell proliferation and differentiation^81^. On the other hand, blocking PDGFRα, which is expressed in fibroblast-like cells has been shown to preserve the BBB^82^. The origin of Col1α1^+^ fibroblast- like cells remains unknown, though some evidence suggests a potential pericyte origin^83^. These cells likely represent a heterogeneous group of fibroblasts, possibly including cells derived from diverse sources such as dural, arachnoid, pial, and perivascular fibroblasts, as well as meningothelial cells with distinct properties and functions in stroke. Recent single-cell transcriptomic studies help us to understand the complex diversity of fibroblasts in developmental and adult brain tissue^84,85^, and future research on stroke tissue could further reveal the unique characteristics of Col1α1+ fibroblast-like cells. Moreover, ICs were identified to express insulin-like growth factor-binding protein 5 (IGFBP5). Additional immunostaining of stroked and intact brain sections revealed upregulation of IGFBP5 in the stroke core, compared to peri-infarct nor the intact tissue. We also found upregulation of IGFBP5 in the analyzed human stroke dataset. IGFBP5, the most conserved member of the IGFBP family, plays various biological roles, such as influencing the inflammatory response^86^, fibrosis^87^, cell adhesion^88^, and cell migration and proliferation^89^. Recent research shows that IGFBP-5 has specific roles depending on the cell type and the physiological or pathological context. It may be involved in the development of atherosclerosis by binding to extracellular matrix (ECM) components PAI-1 and osteopontin, which are found in atherosclerotic plaques and have been shown to promote atherosclerosis in loss-of-function studies^90–92^. *In vitro* studies with primary human idiopathic pulmonary fibrosis (IPF) fibroblasts have shown that both exogenous and endogenously expressed IGFBP-5 increase the expression of ECM component-associated genes and pro-fibrotic genes^93^. Recent findings also suggest that IGFBP5 is essential for regulating angiogenesis. IGFBP5 is induced during reparative angiogenesis in a hind limb ischemia model, and blocking of IGFBP5 has been shown to enhance angiogenesis by boosting ATP metabolism and stabilizing HIF1α via E3 ubiquitin ligase VHL^94^. These results suggest that IGFBP5 could be an interesting pharmacological target for treating conditions related to impaired angiogenesis, such as stroke.^95^

Moreover IGFBP5+ ICs were found to co-express PDGFR- β, further pointing towards a fibroblast-like identity. PDGFRβ-expressing fibroblasts proliferate, migrate, and deposit extracellular matrix at lesion sites after spinal cord injury in mice,^96^ and a similar PDGFRβ^+^/CD105^+^ subset originating from large vessels rather than capillaries contributes to scar formation after ischemic stroke in mice and humans.^97^ These cells do not express pericyte markers NG2 or CD13 but exhibit increased fibronectin expression, strongly supporting a fibroblast-like identity. However, the origins of these fibroblast-like cells and their functions during pathological conditions require future validation.

Our comparative analysis of transcriptomic changes in human and mouse stroke tissue revealed a shared upregulation of genes and pathways, indicative of a conserved response to stroke across species. This analysis has highlighted key genes like CXCL5 and C4B, which are involved in the inflammatory response, suggesting an orchestrated immune activation in the chronic phase of stroke. CXCL5 has been reported as a potential CSF biomarker correlating with brain damage in stroke patients^98^, but also beyond the acute phase patients exhibit a proinflammatory signature after stroke. Elevated CXCL5 proteins have been reported in blood from stroke patients up to 7 years after injury^99^. Another shared pathway was ECM remodeling involving genes such as MMP2 and COL4. Alterations in COL4 expression after stroke have been described in rodent models of MCAO and non-human primates^100,101^. Furthermore, an association between Col4 mutations and ischemic stroke has been described in humans, suggesting that Collagen IV plays an important role in the pathogenesis and recovery of ischemic stroke in both species^102^.

The upregulation of ANGPTL4 suggests the role of angiogenesis in post-stroke recovery. Post-stroke angiogenesis has been reported as an important recovery in experimental and human stroke^30,103–105^. More recently, ANGPTL4 was associated with poor prognosis in acute ischemic stroke patients^106^. The cross-species comparison enhances the validity of these findings and suggests that druggable pathways that show beneficial effects in the mouse may target the same pathway in chronic human stroke.

We recognize that our molecular stroke atlas is only a first step towards deciphering the molecular and cellular interaction following a stroke. We acknowledge limitations such as unintended biases in nuclei isolation and tissue collection from stroke-affected, peri-infarct, and intact tissue, of each mouse. These biases might affect the relative cell proportions and will require future validation using spatially resolved datasets. Another limitation is that, while we have demonstrated the presence of certain interaction partners (e.g., ITGAV and PDGFRβ signaling molecules) at the protein level, further functional studies (e.g., loss-of-function experiments) will be necessary to fully validate the predicted cell-cell interactions and elucidate their roles in post-stroke remodeling. Additional work should extend to validating our findings in longer-term studies, beyond one month, and in alternative stroke models such as the permanent and transient middle cerebral artery occlusion (pMCAo) models in rodents, non-human primates, and human stroke patients. Despite these limitations, our results offer valuable insights for upcoming research into the long-term mechanisms of stroke pathology and the development of therapeutic strategies.

## Materials and Methods

### Experimental design

The study was designed to generate a single-cell atlas of stroke-injured mouse tissue one month following permanent focal cerebral stroke. we used nine stroked adult male and female mice (3-5 months old) with a C57BL/6J background. We validated a successful stroke induction using Laser Doppler imaging and collected tissue from stroke, peri-infarct and intact cortex. We dissociated the tissue and isolated nuclei for subsequent snRNAseq analysis. We compared the mouse transcriptome with a publicly available human stroke dataset (GSE56267).

### Photothrombotic stroke induction

Photothrombotic stroke was induced as previously described ^32,107–110^. Briefly, anesthesia was induced with 4% isoflurane delivered in oxygen. When their respiration rate reached approximately 50 breaths per minute, indicating deep anesthesia, they were placed into a stereotactic frame (Davids Kopf Instruments). A custom-made face mask provided a steady supply of 1-2% isoflurane. Their body temperature was regulated at 36-37 °C using a heating pad. The absence of the toe pinch reflex confirmed deep anesthesia, and Vitamin A eye lubricant from Bausch&Lomb was applied to prevent eye dryness during the procedure. The head was shaved from the neck to the snout, sanitized and Emla™ Creme 5% was applied to the scalp and ears. Ear bars were then inserted to stabilize the head. A 1 cm incision was made to expose the Lambda and Bregma, which were cleaned using a Q-tip. The stroke induction site was precisely marked using an Olympus SZ61 surgery microscope and a WPI UMP3T-1 stereotactic coordinate system, taking Bregma as the reference. Rose Bengal in 0.9% NaCl solution at 15mg/ml was injected intraperitoneally at a dose of 10μl/g bodyweight 5 minutes before a 150W, 3000K Olympus KL1500LCD cold light source was used for illumination at the marked site for 10 minutes. After the procedure, animals were placed in a recovery cage.

### Laser Doppler imaging (LDI)

After stroke induction, anesthetized mice were secured in a stereotactic apparatus. The mice underwent a single-point Laser Doppler Imaging (LDI) procedure using the Moor Instruments MOORLDI2-IR device. LDI data was then extracted and the total flux within the region of interest (ROI) was measured using Fiji (ImageJ). Subsequent analysis was carried out using R software.

### Tissue processing

For RNA sequencing, animals were perfused transcardially on ice using Ringer’s solution (0.9% NaCl). Subsequently, the specified cortical brain tissue was rapidly dissected on ice with the assistance of a microbiopsy instrument (Kai Medical) and a stereotaxic microscope (Olympus). To dissect the core region of the stroke lesion, a micropiopsy punch with a blade diameter of ∅ =1,5 mm was used. To dissect the ischemic border zone, a microbiopsy punch with a diameter of ∅ =2 mm was used. The collected tissue was then immediately frozen in liquid nitrogen. For immunohistochemistry, perfusion was performed using Ringer’s solution (0.9% NaCl), followed by a perfusion with a 4% paraformaldehyde (PFA) solution. The brain was extracted and post-fixed for 6 hours in 4% PFA. The brains were stored in 0.1 M PBS. Before immunohistochemistry, the brains were sectioned into 40 μm coronal slices using a Thermo Scientific HM 450 sliding microtome. Next, brain sections were rinsed with 0.1 M PBS. They were then treated with 500 µl of blocking buffer (5% donkey serum in 1x PBS with 0.1% Triton® X-100) and incubated for 1 hour at room temperature. Following blocking, the sections were incubated with primary antibodies (Table 1) on an Unimax 1010 shaker set to approximately 90 rpm, and this was maintained overnight at 4°C. The next day, after washing, the sections were incubated with appropriate secondary antibodies (Table 2), for 2 hours at room temperature. Additionally, the sections were treated with DAPI (Sigma, diluted 1:2000 in 0.1 M PBS) to stain the nuclei. Finally, the sections were arranged on Superfrost Plus™ microscope slides, immersed in Mowiol® mounting medium, and securely coverslipped.

**Table 1:** Primary Antibody List.

| Antigen | Target | Host | Dilution | Company |
| --- | --- | --- | --- | --- |
| CD13 | Pericytes | goat | 1:200 | R&D Systems |
| CD31 | Vascular endothelial cells | rat | 1:50 | BD Biosciences |
| GFAP | Astrocytes | mouse | 1:200 | R&D Systems |
| Iba1 | Microglia | goat | 1:200 | R&D Systems |
| CD68 | Macrophages | rat | 1:200 | BioLegend |
| vGlut1 | Glutamate transporter in synaptic vesicles | mouse | 1:300 | Synaptic Systems |
| GAT1 | GABA transporter in synaptic vesicles | rabbit | 1:200 | Abcam |
| IGFBP5 | Insulin-like growth factor-binding protein 5 | rabbit | 1:100 | Proteintech |
| NeuN | Neurons | rat | 1:200 | Abcam |

**Table 2:**
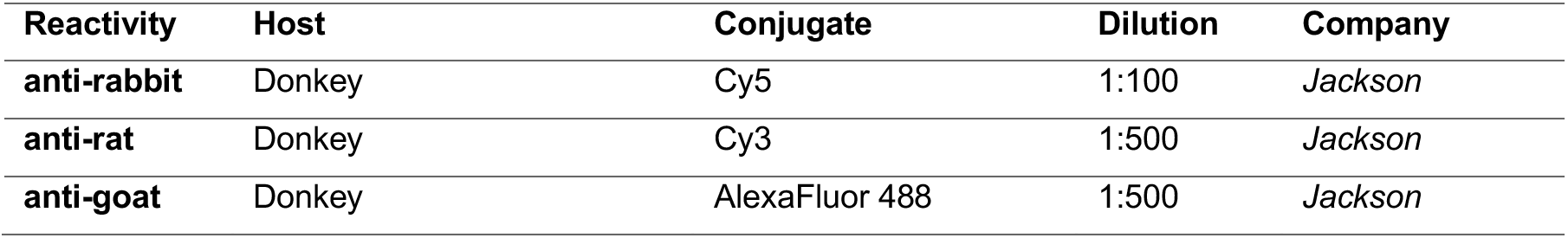
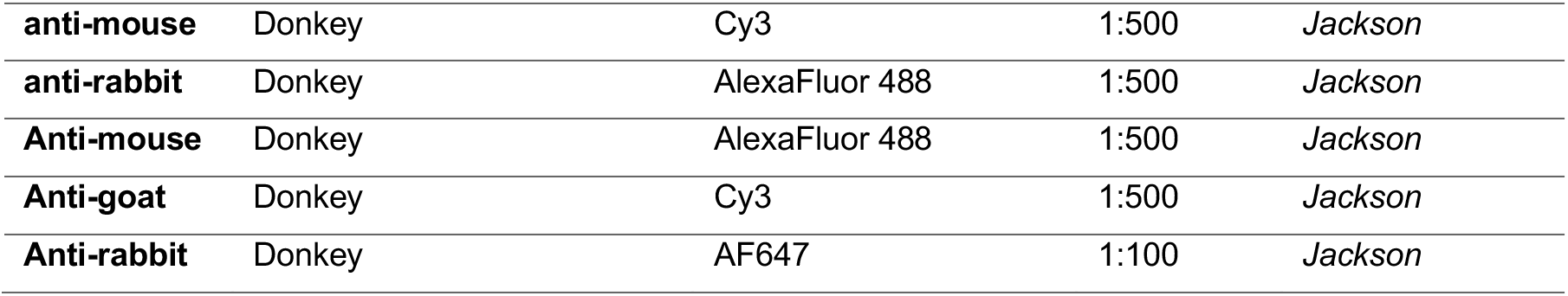
Secondary Antibody List.

| Reactivity | Host | Conjugate | Dilution | Company |
| --- | --- | --- | --- | --- |
| anti-rabbit | Donkey | Cy5 | 1:100 | Jackson |
| anti-rat | Donkey | Cy3 | 1:500 | Jackson |
| anti-goat | Donkey | AlexaFluor 488 | 1:500 | Jackson |
| **anti-mouse** | Donkey | Cy3 | 1:500 | *Jackson* |
| **anti-rabbit** | Donkey | AlexaFluor 488 | 1:500 | *Jackson* |
| **Anti-mouse** | Donkey | AlexaFluor 488 | 1:500 | *Jackson* |
| **Anti-goat** | Donkey | Cy3 | 1:500 | *Jackson* |
| **Anti-rabbit** | Donkey | AF647 | 1:100 | *Jackson* |

### Histological quantification of vasculature, microglia and astrocytes

All analysis steps were conducted on 40 μm coronal sections. The ischemic area was visually identified as defined area with atypical and atrophic tissue morphology including pale areas, the peri-infarct region adjacent to the stroke core was referred to as the ischemic border zone (ibz), extending up to 300 μm around the clearly defined lesion boundary. Cell counting and analysis of fluorescence intensity was assessed using the software FIJI (ImageJ, version 2.1.0/1.53c). For cell counting, we used the built-in plugin “cell counter”. For signal intensity measurements, images were converted to 8-bit format and subjected to thresholding using mean grey values obtained from regions of interest (ROIs) in the unaffected contralateral cortex to create a binary image. Signal intensity was then measured in the respective ROIs using the polygon tool.

### Isolation of nuclei from frozen brains

Nuclei were extracted from frozen cortical brain tissues as previously described^111^. Tissues were rapidly frozen in liquid nitrogen and subsequently pulverized using a Dounce homogenizer in a lysis solution composed of 10 mM Tris-HCl (pH 7.5), 10 mM NaCl, 3 mM MgCl2, and 0.1% Nonidet P40 dissolved in nuclease-free water. After a 15-minute incubation, the homogenate was strained through a 30 µm cell strainer. The strained suspension was then subjected to a low-speed centrifugation at 500g for 5 minutes at 4°C to sediment the nuclei. These nuclei were washed and passed through a 40 µm cell strainer twice, using sterile PBS supplemented with 2% BSA and 0.2 U/µl RNase inhibitor. The nuclei were then suspended in 500 µl of the washing buffer and combined with 900 µl of 1.8 M sucrose solution. This mixture was carefully overlaid onto 500 µl of 1.8 M sucrose and centrifuged at 13,000g for 45 minutes at 4°C, facilitating myelin removal. The final pellet was resuspended in the washing buffer and filtered once more through a 40 µm cell strainer to ensure purity.

### Single-nucleus RNA sequencing

Single-nucleus RNA sequencing (snRNAseq) was performed as previously described.^111^ For the droplet-based library construction, nuclei isolated from both stroked and non-stroked mouse cortices were processed on the Chromium system by 10x Genomics, following the manufacture’s guidelines. RNA capture and subsequent amplification were facilitated by the Chromium Single Cell 3’ Reagent Kits v3. Sequencing of the resulting libraries was executed on an Illumina sequencing platform. The analysis pipeline, including demultiplexing of samples, processing of barcodes, and enumeration of single cells, was conducted using the Cell Ranger Single-Cell Software Suite supplied by 10x Genomics.

### Clustering and annotation cell types

For the clustering and annotation of cell types, single-nucleus RNA sequencing (snRNAseq) data alignment and gene quantification were performed using Cellranger v3.1.0, which followed default settings and referenced the mm10 2020-A dataset. Cells were screened, excluding any with greater than 5% mitochondrial gene expression or with less than 500 nFeature_RNA. Normalization and scaling of gene counts were performed using Seurat v5.0.192, to adjust for total unique molecular identifier counts per cell. Using the initial 30 principal components, cell clustering was accomplished via the FindNeighbors function, with subsequent clustering by the FindClusters function. Dimensionality was reduced through Uniform Manifold Approximation and Projection (UMAP) employing the RunUMAP function. Distinct cell types were identified based on established markers, categorizing into Glutamatergic neurons (Glut: e.g., Slc17a7, Satb2), GABAergic neurons (GABA: e.g., Gad1, Gad2), Astrocytes (Asc: Slc1a2, Slc1a3), Fibroblasts (FB: Col1a, Fn1), Oligodendrocytes (Olig: Mbp, Plp1), Immune cells (IC: Inpp5d, Csf1r), vascular cells (Vasc: Flt1, Cldn5), stem and progenitor cells (SPC: Sox10, Vcan), and mural cells (Per: Pdgfrb Cspg4). The cell types and marker expression of cell types matched sc/snRNAseq datasets^12,26,27^. FindMarkers function was used to pinpoint cell type–specific marker genes, considering genes with a Bonferroni correction adjusted P value <0.05 as significant markers. The downstream analysis (including Gene Set Enrichment analysis (GSEA), and over representation analysis (ORA) was performed using R package ClusterProfiler.

### Cell-cell communication with CellChat

Differential cell-cell interaction networks were generated using CellChat version 2.1.0^47,48^. In brief, DifferentialConnectome was applied to the Seurat objects (version 5.01), which contained integrated data of mouse stroke, ibz and intact datasets. The compareInteractions function was utilized to compute the total number of interactions and their strengths, while network centrality was scored using the netAnalysis_computeCentrality function. All analyses were conducted in line with the ’Full tutorial for CellChat analysis of a single dataset with detailed explanation of each function’ found on the GitHub page.

### Human RNAseq dataset

Human stroke RNA sequencing data was obtained from the NCBI Gene Expression Omnibus, accession number GSE56267. For the purpose of comparing differential gene expression profiles between mouse and human stroke, we employed a pseudo-bulk approach on the mouse snRNAseq data using the AggregateExpression() function in the Seurat package. We processed both datasets using Z-score normalization and translated mouse gene identifiers to their human orthologs. To evaluate the relationship between gene expression in mouse and human stroke cases, Pearson’s correlation test was used. Subsequent analyses, which included Gene Set Enrichment Analysis (GSEA) and Over Representation Analysis (ORA), were conducted using the ClusterProfiler package in R, providing comprehensive insights into the biological significance of the expression data.

### Statistical analysis

Statistical analysis was performed using RStudio (Version 4.04). Sample sizes were designed with adequate power in line with previous studies from our group^112–114^ and relevant literature. One-way analyses of variance (ANOVA) followed by Tukey multiple comparison test was performed for cerebral blood flow measurements. The assumption of normality was tested by Kolmogorov–Smirnov tests and by inspecting residuals with QQ plots. Data is expressed as mean ±SD; statistical significance was defined as *p < 0.05, **p < 0.01, and ***p < 0.001.

### Data availability

Raw single nucleus RNA sequencing data is deposited in the NCBI Gene Expression Omnibus (GEO) under the accession number GSE276202. Data are now also available to explore via an interactive web browser: https://rustlab.shinyapps.io/Stroke-Atlas/

## Ethics declaration

All in vivo experiments were performed at the Laboratory Animal Services Center (LASC) in Schlieren, Switzerland according to the local guidelines for animal experiments and were approved by the Veterinary Office of the Canton Zurich in Switzerland (Protocol number 209/2019).

## Acknowledgement

This work is supported by the Swiss 3R Competence Center (OC-2020-002) and the Swiss National Science Foundation (CRSK-3_195902) and (PZ00P3_216225) to RR. The work was also supported by the National Institutes of Health (R01NS117827) and by the development funds to the Center for Neurodegeneration and Regeneration at the Zilkha Neurogenetic Institute. In addition, RR and CT acknowledge support from the Mäxi Foundation.

## Author contribution

RZW, CT, RR contributed to overall project design. RZW, KK, RR contributed to the design of snRNAseq experiments. RZW, BAB, NHR, RR conducted and analyzed *in vivo* experiments. RZW, AB, MZ, RR performed nuclei isolation and snRNAseq experiments. RR analyzed snRNAseq experiments. RZW, RR made figures. CT, RR supervised the study. RZW, KK, CT, RR wrote and edited the manuscript with input from all authors. All authors read and approved the final manuscript.

