## Supplementary material for "A molecular brain atlas reveals cellular shifts during the repair phase of stroke": SI

### Correspondence

**Ruslan Rust, PhD**

Assistant Professor
The Zilkha Neurogenetic Institute
Department of Physiology and Neuroscience
Keck School of Medicine of the University of Southern California
1501 San Pablo Street, Room 341
Los Angeles, CA 90033

X: @rust_ruslan
ORCID: 0000-0003-3376-3453

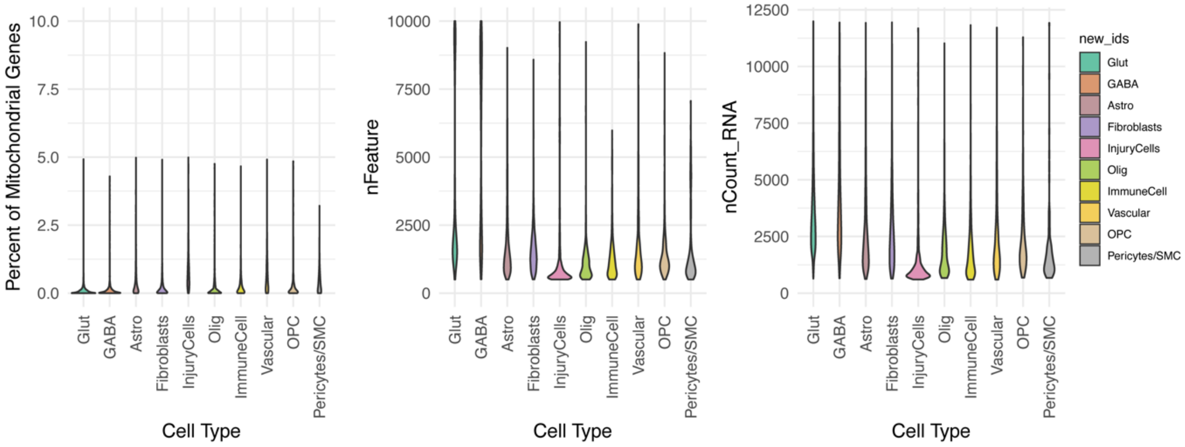


**Suppl. Fig. 1. Quality control assessment of pre-processed single-cell RNA-seq datasets**. Violin plots display the distribution of mitochondrial gene expression percentage (left), number of detected features (nFeature, center), and total RNA counts (nCount, right) across different cell types.

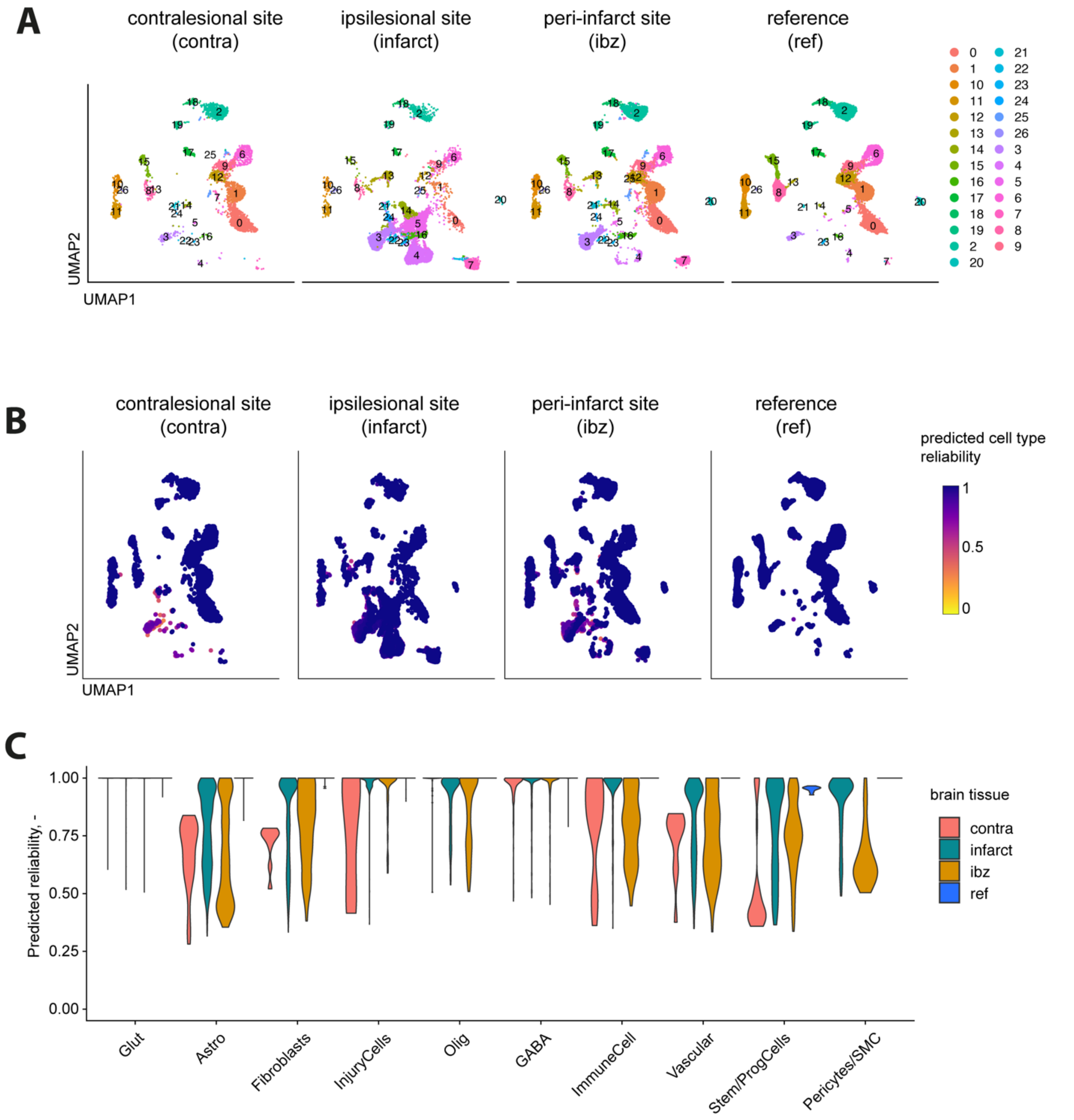


**Suppl. Fig. 2: Identification and validation of cell annotations across different brain regions one month after stroke.** (A) UMAP of initial Seurat clustering for the contralesional site (contra), ipsilesional site (infarct), peri-infarct site (ibz), and reference mouse cortex (ref). (B) UMAP showing predicted cell type reliability across all datasets, with color intensity reflecting the confidence level in cell type assignment (from low in yellow to high in dark blue). (C) Violin plot displaying the predicted reliability of cell type annotations across different brain tissues and cell types.

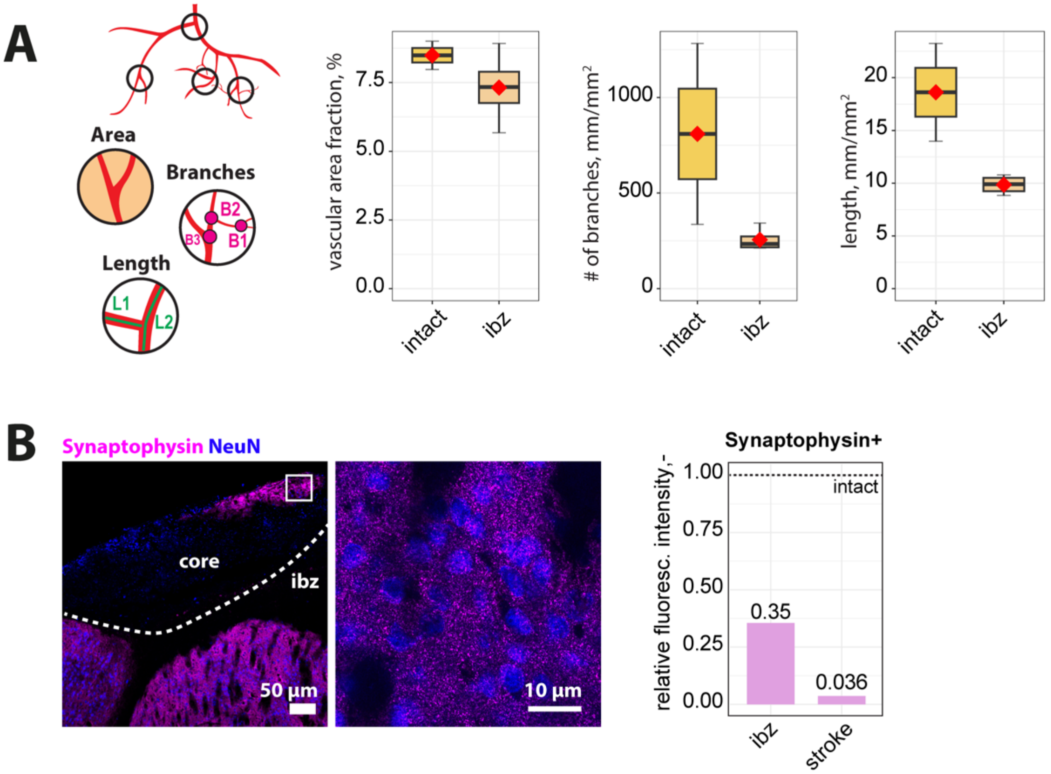


**Suppl. Fig. 3: Vascular and synaptic changes after stroke.** (A) Quantification of vascular density, including area, branch number, and length, in the intact brain and the ischemic border zone (ibz) one month after stroke. (B) Representative histological overview of brain sections stained with synaptophysin and NeuN and quantification of synaptophysin fluorescence intensity in ischemic border zone (ibz) and stroke core areas one month after stroke.

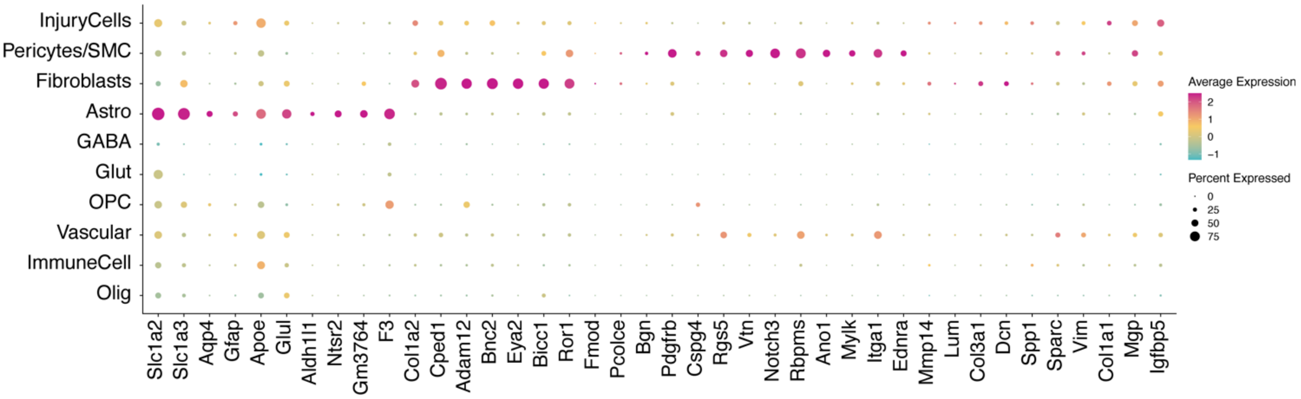


**Suppl. Fig. 4: Cell type marker for astrocytes, fibroblasts, pericytes and injury-associated cells are distinct in stroke tissue.** Dot plot representation of cell type markers across different cell populations labeled by cell type: glutamatergic neurons (Glut), GABAergic neurons (GABA), astrocytes (Asc), fibroblasts (FB), oligodendrocytes (Olig), immune cells (Imm), vascular cells (Vasc), stem/progenitor cells (SPC), and mural cells (Per)

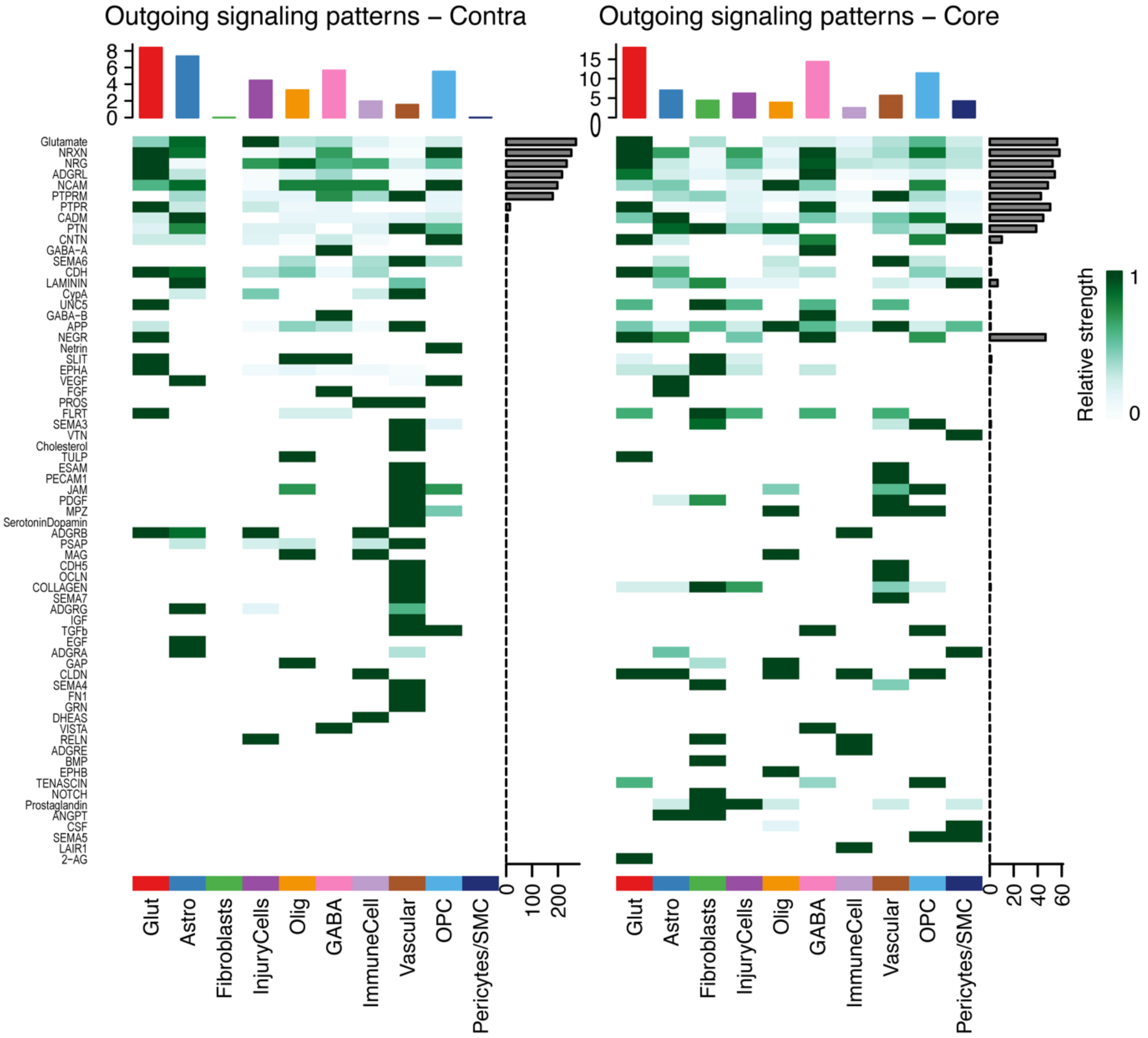


**Suppl. Fig. 5: Predicted outgoing signals during cell-cell communication.** Pathways that contribute to sending signals to individual cell types in contralesional (contra, left) and stroke (core, right) tissue one month after stroke.

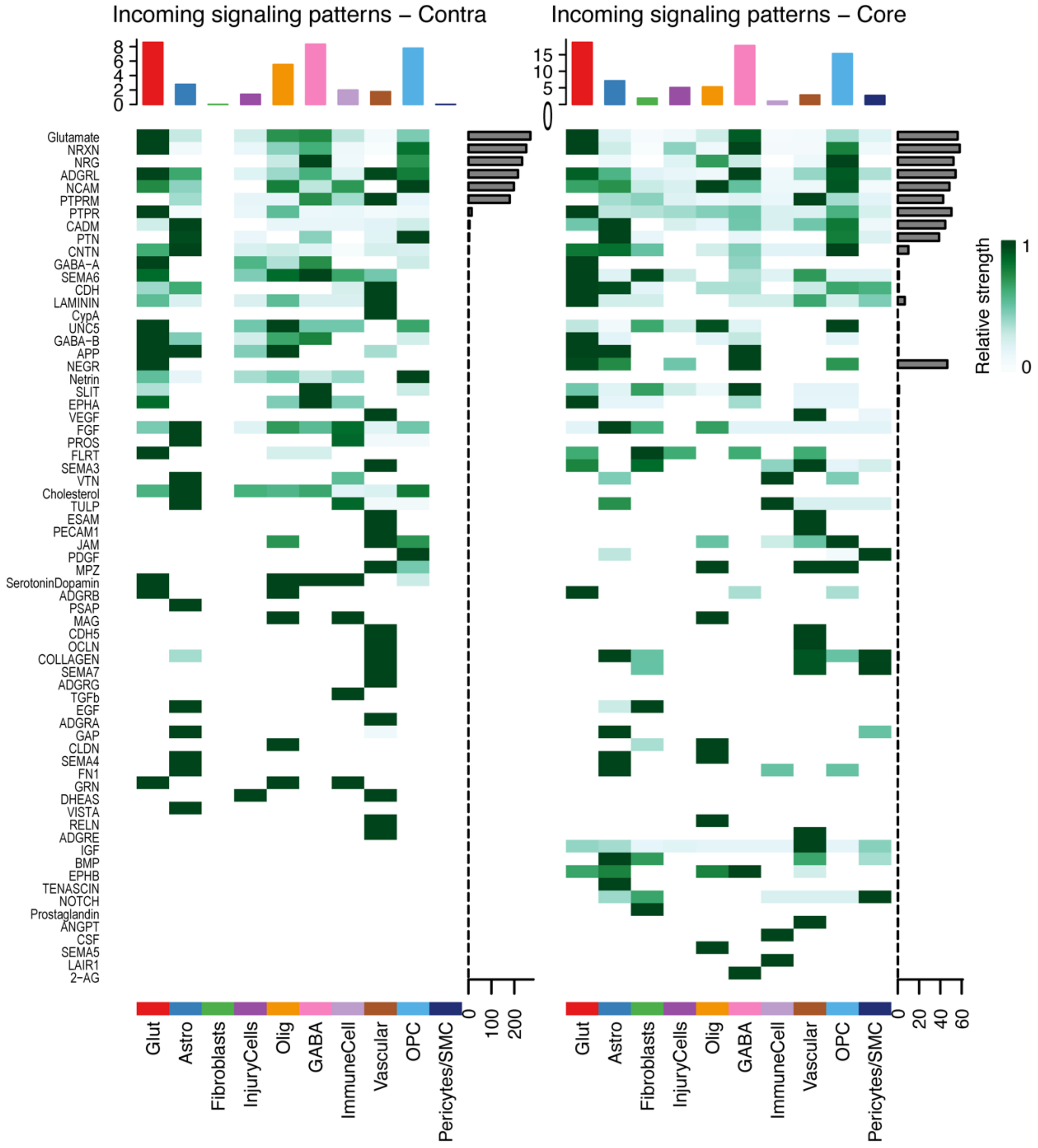


**Suppl. Fig. 6: Predicted incoming signals during cell-cell communication.** Pathways that contribute to receiving signals from individual cell types in contralesional (contra, left) and stroke (core, right) tissue one month after stroke.

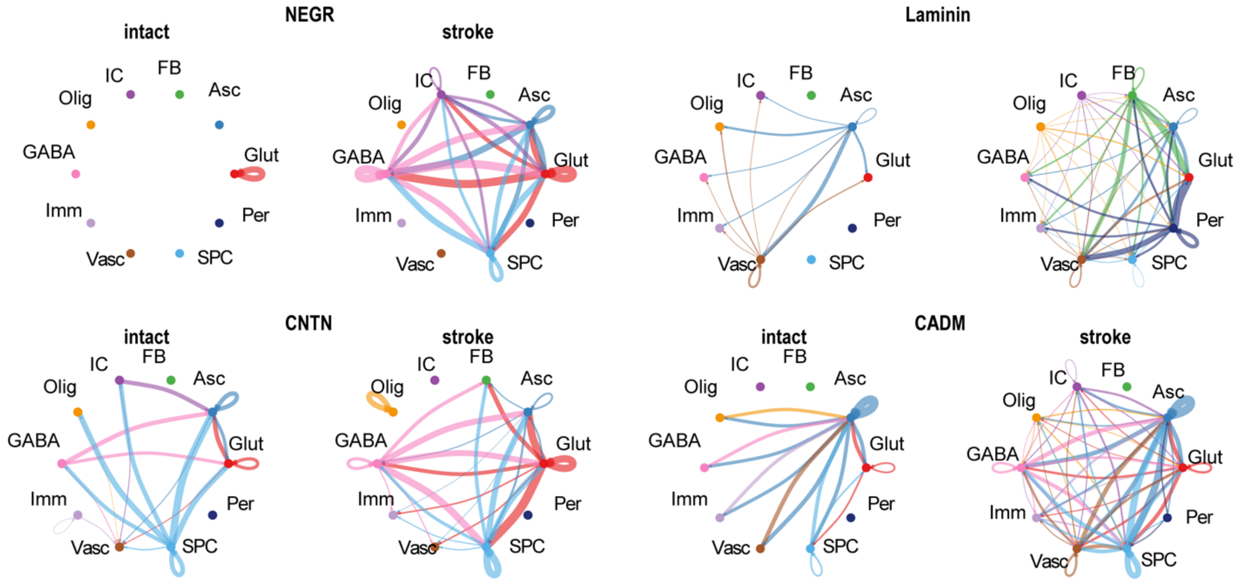


**Suppl. Fig. 7: Analysis of specific communication pathways in intact and stroke tissue**. Interactions are shown from each cell types that are increased in stroke compared to intact tissue for the pathways involving neural growth regulator (NEGR), Laminin, Contactin (CNTN) and Cell adhesion molecules (CADM) one month after stroke.
